# Routes to Roots: Direct Evidence of Water Transport by Arbuscular Mycorrhizal Fungi to Host Plants

**DOI:** 10.1101/2020.09.21.305409

**Authors:** Anne Kakouridis, John A. Hagen, Megan P. Kan, Stefania Mambelli, Lewis J. Feldman, Donald J. Herman, Jennifer Pett-Ridge, Mary K. Firestone

## Abstract

Arbuscular mycorrhizal fungi (AMF) form symbiotic associations with 80% of surveyed land plant species and are well-recognized for accessing and transferring nutrients to plants^1^. Yet AMF also perform other essential functions, notably improving plant-water relations^2^. Some research attributes the role of AMF in plant-water relations solely to enhancing plant nutrition and osmoregulation for plants partnered with AMF^3,4,5^, while indirect evidence suggests AMF may transport water to plants^1,6,7^. Here, we used isotopically-labeled water and a fluorescent dye to directly track and quantify water transport by AMF to plants in a greenhouse experiment. We specifically assessed whether AMF can access water in soil unavailable to plants and transport it across an air gap to host plants. Plants grown with AMF that had access to a physically separated ^18^O-labeled water source transpired twice as much, and this transpired water contained three times as much label compared to plants with AMF with no access to the separated labeled water source. We estimated that water transported by AMF could explain 46.2% of the water transpired. In addition, a fluorescent dye indicated that water was transported via an extracytoplasmic hyphal pathway.

---

Water availability limits plant growth and is an ever-pressing issue in the context of climate change^8^. Plants have evolved multiple strategies to increase their tolerance of water deficit and alleviate its detrimental effects^9^, including associations with AMF.

AMF can improve plant-water relations via several indirect mechanisms^2^. Specifically, AMF help regulate stomatal conductance^3,10,11^ and hydraulic properties of roots^12,13^, improve plants’ ability to osmoregulate^5,14^, and reduce drought-induced oxidative stress in their host plants^4^.

A small number of studies have suggested that AMF may transport water to their host. While investigating nutrient transport, Faber *et al*. (1991) discovered that plants with intact AMF hyphae transpired 37% more than plants with severed hyphae. Khalvati *et al*. (2005) found that when AMF were allowed to access a separate compartment from which roots were excluded, the compartment weighed 4% less at the end of the experiment.

The potential pathways AMF hyphae use to move water to roots are unknown. Water could be transferred via hyphae by travelling along the outside of the fungal cell wall or through the cell wall matrix itself^15^. We refer to this potential pathway as ‘extracytoplasmic’, in contrast to a ‘cytoplasmic’ pathway where the cell-to-cell transport occurs inside the fungal cell membrane. These terms are analogous, but not identical, to what is traditionally described as apoplastic and symplastic transport in plants.

To resolve this knowledge gap, we investigated if and how the AMF *Rhizophagus intraradices* transports water to the host plant *Avena barbata*, wild oats. *R. intraradices* naturally colonizes *A. barbata* roots at our annual grassland field site near Hopland, CA, where *A. barbata* seeds were collected. In a greenhouse study, three *A. barbata* seedlings were planted in a compartment filled with a sand-clay mixture, the ‘plant compartment’, of two-compartment microcosms (fig. 1 and supplementary material fig. 1). The other compartment, the ‘no-plant compartment’, was filled with a soil-sand mixture and was separated from the plant compartment by a 3.2 mm air gap to prevent liquid water from traveling via mass flow between compartments. Each side of the air gap was covered by nylon mesh. Mesh (18 μm) that allowed hyphae but excluded roots was used for AMF-permitted, H_2_^18^O enriched water/dye microcosms, termed ‘+AMF’ (treatment 1), and AMF-permitted natural abundance water/no dye controls, termed ‘^16^O’ (treatment 2). Mesh (0.45 μm) that excluded both hyphae and roots was used for AMF-excluded H_2_^18^O enriched water/dye controls, termed ‘-AMF’ (treatment 3). For all three treatments, plants were inoculated with *R. intraradices* and well-watered for the first three weeks, then kept under low water for an additional seven weeks (approximate mean gravimetric water content = 16.9%, mean water potential = −1.5 MPa). The plant and no-plant compartments of all microcosms received a nutrient solution weekly, and -AMF plants received twice as much to counter the nutrients +AMF plants had access to via AMF in the no-plant compartment. Once plants were fully-grown, we injected ^18^O-labeled water with a fluorescent dye into the no-plant compartment of the +AMF and -AMF microcosms. Unenriched water that did not contain the fluorescent dye was injected into the ^16^O microcosms to assess natural isotope fractionation and to provide a control for root autofluorescence. We collected water transpired by plants for three days and then harvested plants, hyphae, sand-clay and soil-sand mixtures.

**Figure 1:**
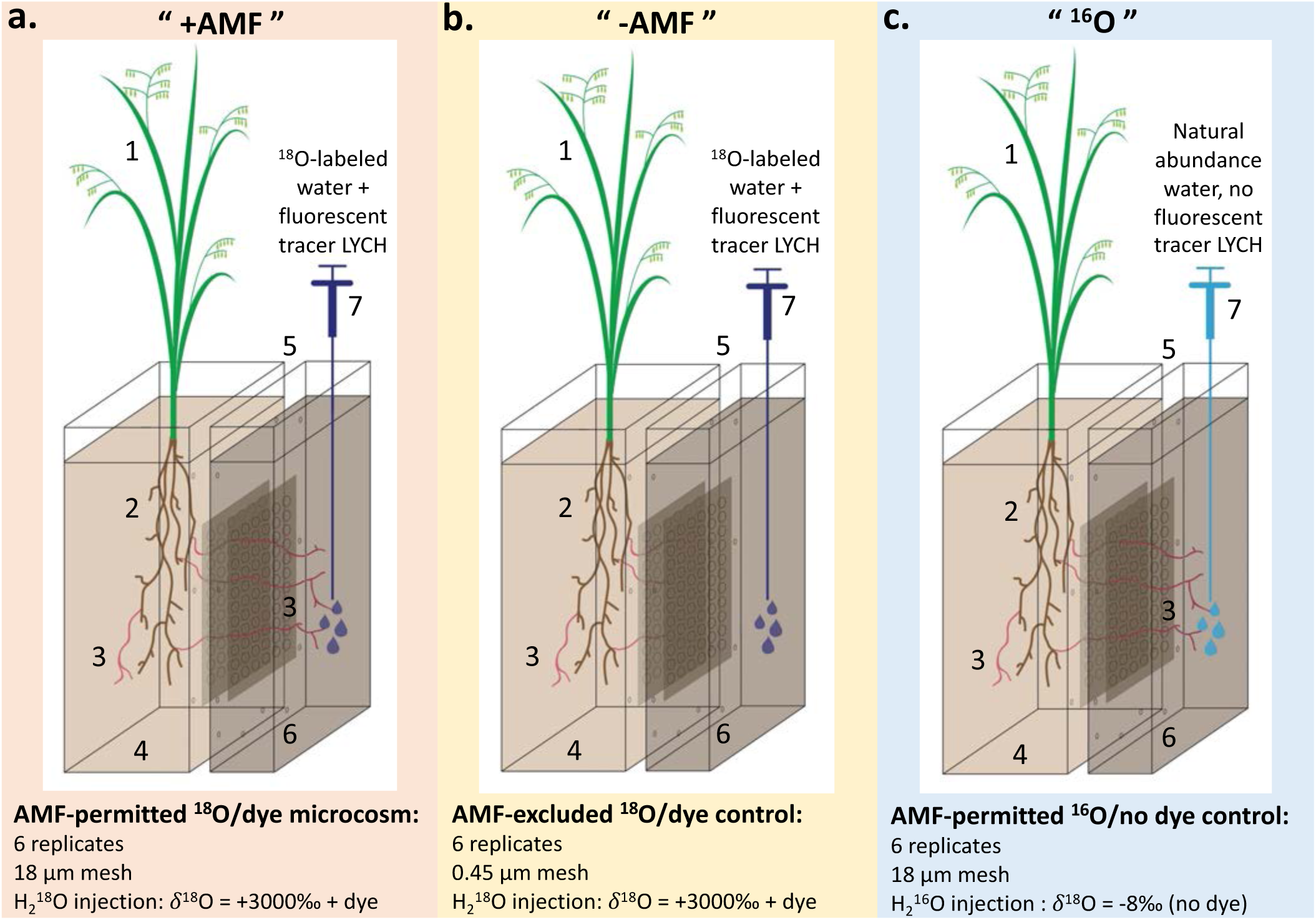
Experimental set up designed to test the movement of water to plants by AMF hyphae. **a. AMF permitted** ^**18**^**O/dye microcosm (“+AMF”)** where AMF are able to access a no-plant compartment, with ^18^O-labeled water and fluorescent tracer LYCH injected into the no-plant compartment. **b. AMF excluded** ^**18**^**O/dby**.**e control (“-AMF”)** where AMF are not able to access the no-plant compartment, with ^18^O-labeled water and fluorescent tracer LYCH injected into the no-plant compartment. **c. AMF permitted** ^**16**^**O/no dye control (”**^**16**^**O”)** where AMF are able to access the no-plant compartment and natural abundance water without a fluorescent tracer is injected into the no-plant side. In **a**.**-c**.**: 1**. *Avena barbata* shoots **2**. *A. barbata* roots **3**. AMF *Rhizophagus intraradices* **4**. Plant compartment filled with 1/2 sand 1/2 clay mixture **5**. 3.2 mm air gap. **6**. No-plant compartment filled with 1/2 soil 1/2 sand mixture **7**. Syringe illustration of injection of solutions into the no-plant compartment.

Using Sanger sequencing of roots, hyphae, and soil-sand mixture, we confirmed that *R. intraradices* colonized roots in all microcosms and grew hyphae across the air gap and mycelium into the no-plant compartment in +AMF and ^16^O microcosms. Using light and fluorescence microscopy, we confirmed that *R. intraradices* was active in roots by staining them with acid fuchsin and observing hyphae, spores, and arbuscules (fig. 2.a.-d. and supplementary material fig. 2). We also observed hyphae crossing the air gap, and extensive hyphal networks in the soil-sand mixture of the no-plant compartment of +AMF and ^16^O microcosms (fig. 2.e.-f.). In the no-plant compartment of the -AMF microcosms, we did not observe hyphae crossing the air gap nor hyphal networks in the soil-sand mixture.

**Figure 2:**
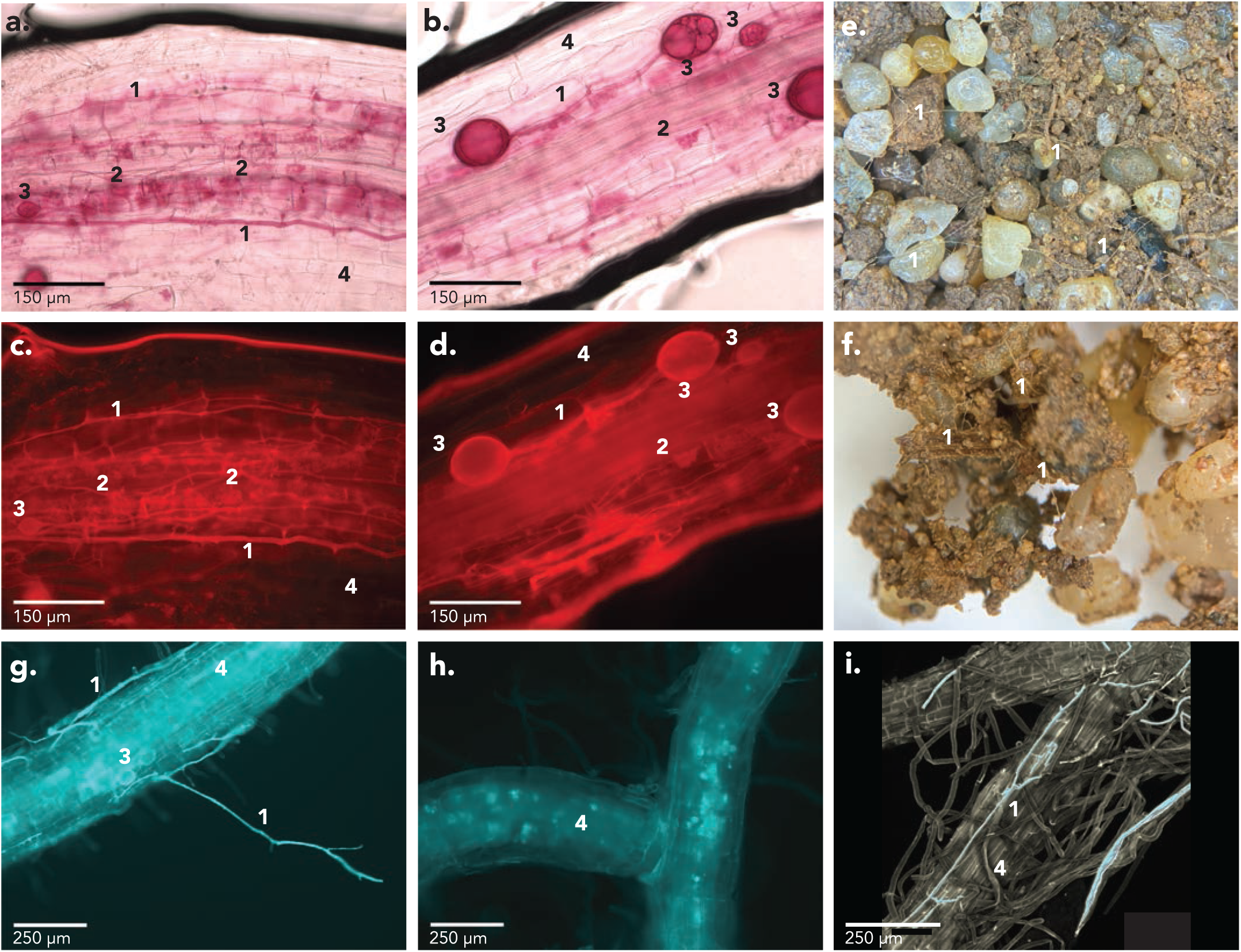
**a-d**. *Avena barbata* roots dyed with acid fuchsin showing AMF structures. **a&b**. Bright field micrographs. **c&d**. Fluorescence images at AMF wavelengths (**λ**ex 596 nm; **λ**em 615 nm). **e&f**. Soil-sand mixture from the no-plant compartment of a +AMF microcosm with numerous AMF hyphae visible under a dissecting microscope. **g-i**. Fluorescence micrographs of roots at LYCH wavelengths (**λ**ex 428 nm; **λ**em 536 nm). **g**. Root from a +AMF microcosm with hyphae and vesicles visible in blue. h. Root from a ^16^O control microcosm where hyphae and vesicles not visible, only root autofluorescence. i. Reconstituted 3D model from confocal images of a root from a +AMF microcosm; fluorescing tissues are blue, non-fluorescing tissues are grey. In **a-i**.: **1**. Hyphae. **2**. Arbuscules. **3**. Vesicles **4**. Root.

We found that +AMF plants transpired almost twice as much water as -AMF plants, 7.80 and 4.02 mL respectively over three days (*P* < 0.05) (fig. 3.a. and supplementary material table 1). The gravimetric water content (and water potential) in the plant and no-plant compartments, the mass of above-ground biomass (a proxy for leaf area index), and the root:shoot ratio, were not significantly different between treatments (supplementary material table 1). We observed no difference in the carbon to nitrogen ratio (C:N) and percent nitrogen (N) of the plant shoots between treatments. General plant stunting, a red/purple color, and tissue nutrient content suggested that plant biomass was limited by phosphorus (P) availability for both +AMF and - AMF plants; -AMF plants had a significantly lower P content than +AMF plants (*P* < 0.05, supplementary material table 1). The excess water transpired by +AMF plants is likely due to the presence of AMF hyphae accessing water in the no-plant compartment.

**Figure 3.**
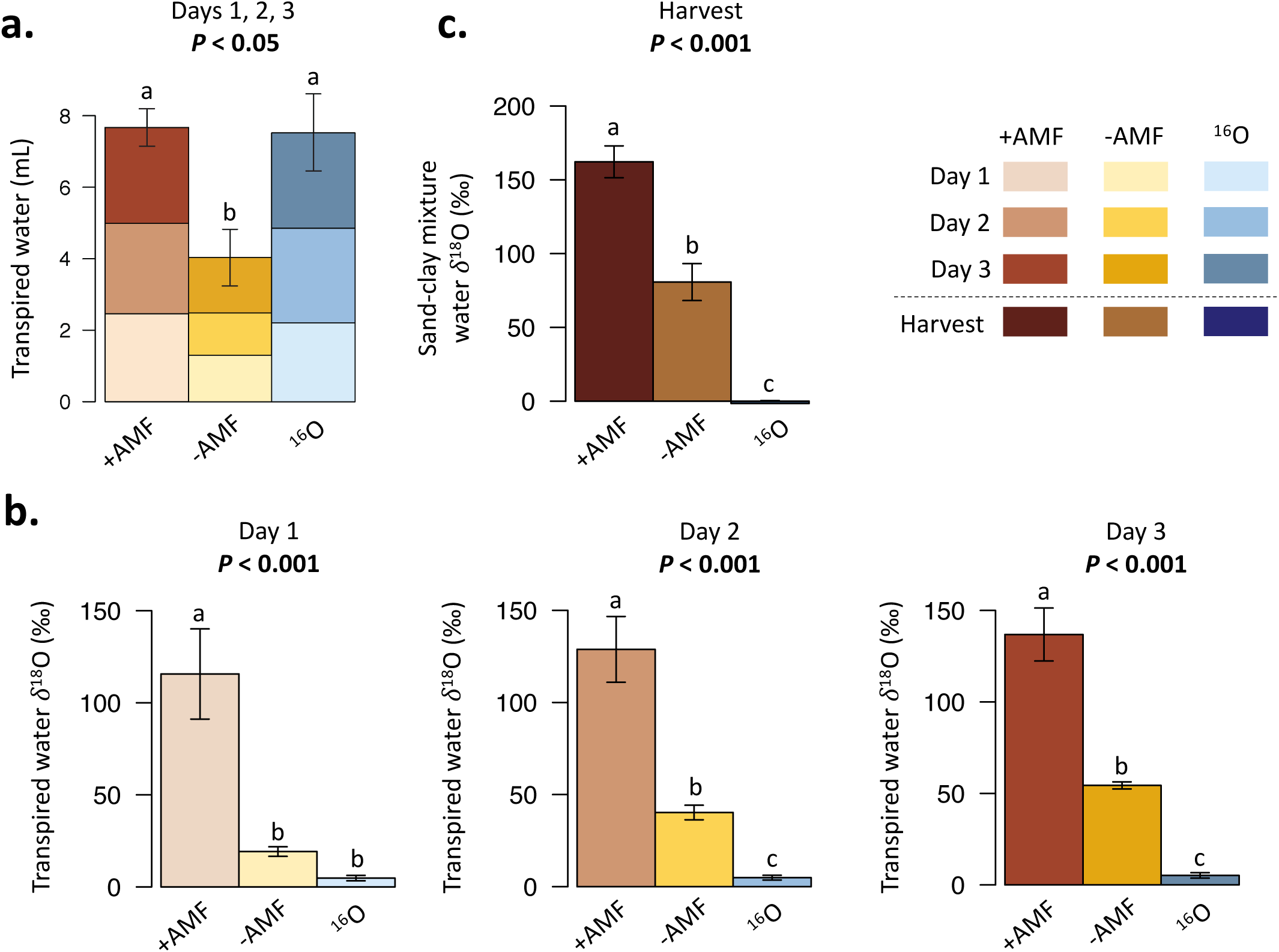
Volume and isotope enrichment of water transpired by *Avena barbata* shoots over three days in +AMF, -AMF, and ^16^O microcosms. Each color and shade (light, medium, dark) represents one day of water transpired. **a**. Volume of water transpired. **b**. δ^18^O value of transpired water. **c**. δ^18^O value of water from the sand-clay mixture in the plant compartment after destructive harvest of the microcosms. In a, b, c, different letters above bars represent statistically significant differences (one-way ANOVA & Fisher LSD test); corresponding *P*-values are indicated above each plot. The error bars represent standard error (n=6).

To test whether +AMF plants obtained water from the no-plant compartment via AMF, we quantified the ^18^O enrichment of transpired water. On average, transpired water from +AMF plants was three times as ^18^O enriched relative to that from -AMF plants (δ^18^O 127.09‰ and 41.28‰, respectively, *P* < 0.001) (fig 3.b. and supplementary material table 1). The water in the sand-clay mixture in the plant compartment of the +AMF microcosms was twice as ^18^O enriched relative to -AMF microcosms (δ^18^O 162.19‰ and 80.7‰, respectively, *P* < 0.001) (fig. 3.c. and supplementary material table 1). While -AMF controls also had a higher δ^18^O in transpired water and sand-clay mixture water than ^16^O controls (presumably some of the ^18^O water from the no-plant compartment travelled passively via water vapor or droplets in the air gap into the plant compartment), there was significantly more ^18^O in transpired water and sand-clay mixture water in +AMF microcosms, implying that more ^18^O-labeled water travelled via hyphae than by other methods. These data allow us to directly confirm that excess water transpired by +AMF plants came from the no-plant compartment via AMF. These results, to our knowledge, are the first direct evidence of water transport by AMF to host plants.

Plants in +AMF microcosms transpired an average of 7.80 mL of water over three days. Using the measured volumes and δ^18^O of transpired water in +AMF and -AMF microcosms (supplementary material table 1), we estimated that the total AMF-transported water amounts to an average of 3.60 mL over three days per microcosm, each holding three plants. Thus, AMF-transported water accounts for 46.2% of the total transpired water by +AMF plants (detailed calculations can be found in the Methods section).

Finally, we used a fluorescent tracer to test the specific pathway of water transport by fungal hyphae. The ^18^O-labeled water we injected into the no-plant compartment of +AMF and - AMF microcosms also contained the membrane-impermeant fluorescent dye lucifer yellow carbohydrazide (LYCH). LYCH can travel on the outer surface of the hyphal cell wall and inside the hyphal cell wall matrix, but it cannot cross cell membranes into the cytoplasm^16^. This property allowed us to investigate hyphal extracytoplasmic transport. In figure 2 g.-h., this dye can be seen on hyphae outside and inside roots, and in vesicle cell walls inside roots, indicating that it was travelling via extracytoplasmic hyphal transport in +AMF microcosms. LYCH has a high affinity for the cell wall matrix^17^ and did not diffuse out once taken up by hyphae. In both - AMF and ^16^O microcosms, no hyphae or spores are fluorescing; only naturally occurring root autofluorescence at these wavelengths^18^ can be observed (fig 2. h.-i.).

In plants, roots can transport water via both symplastic and apoplastic pathways, and plants can regulate the relative contribution of each route based on environmental conditions^19^. The symplastic pathway, which tends to be favored when water availability is limited^20,21^, is slower because water has to flow from cell to cell via the cytoplasm, crossing plasma membranes or plasmodesmata, following an osmotic gradient^19,20^. The apoplastic pathway, which is favored when plants are not water-stressed^20,21^, is faster because water travels extracellularly through the cell wall and matrix and moves directly and continuously via the transpiration stream, facing little resistance^20^. Interestingly, Barzana *et al*. (2012) found that plants with AMF associations have an increased apoplastic water flow in both drought and non-drought conditions, and have a greater ability to switch between water transport pathways, compared to plants with no AMF. They further suggest that AMF hyphae could contribute water to the apoplastic flow in roots, consistent with our observations. It appears that AMF can act as extensions of the root evapotranspiration pathway, with plant transpiration driving water flow along hyphae outside of the hyphal cell membrane.

Our study provides strong evidence to support the existence of extracytoplasmic transport in hyphae. It is possible that cytoplasmic transport could occur at the same time. AMF and plants both have aquaporins at the arbuscule-plant cell interface^12,13,22,23^, and AMF have been shown to increase root hydraulic conductivity and symplastic flow in roots under drought conditions^12,13^. In our experiment, we used a second fluorescent tracer, one that can cross cell membranes, to track water moving via cytoplasmic transport from hyphae to plants. However, this tracer diffused out of hyphae, so it was not possible to distinguish whether the dye actually moved within the hyphal cytoplasm versus outside the membrane (data not included). A summary of our proposed model of water flow from hyphae to roots is shown in figure 4.

**Figure 4:**
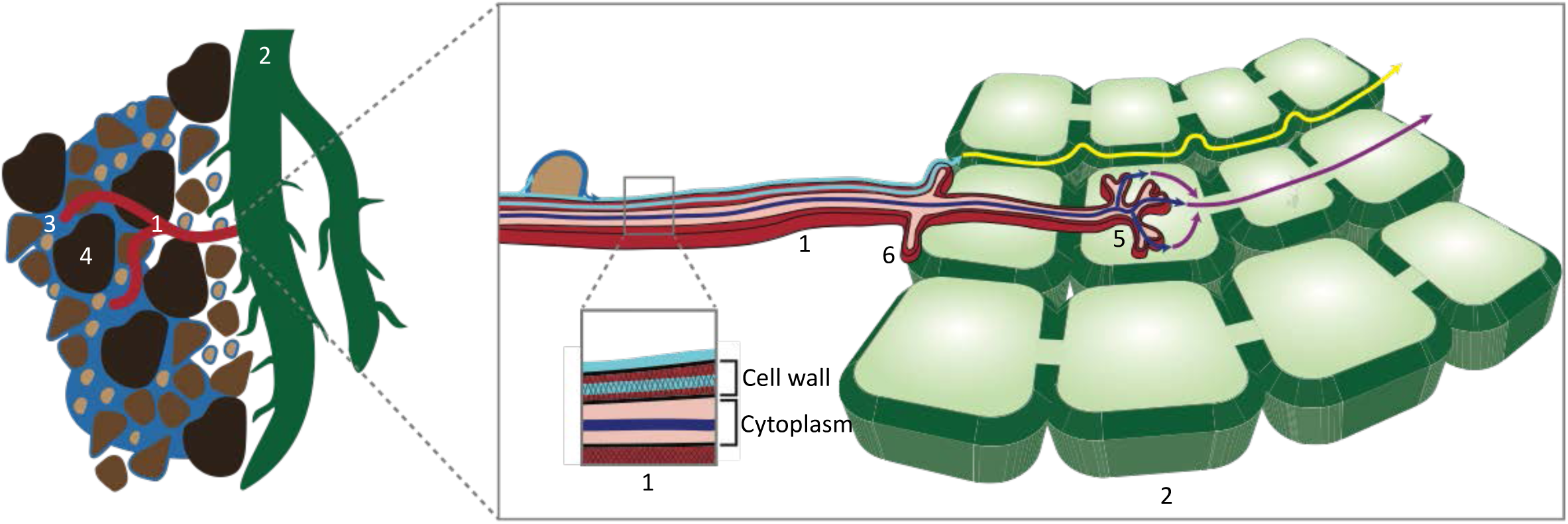
Simplified representation of water transport from soil through AMF hypha to a plant root. **Extracytoplasmic water transport** in a hypha, represented by a light blue arrow, joins **apoplastic transport** in a plant root, represented by a yellow arrow. **Cytoplasmic transport** in a hypha, represented by a dark blue arrow, joins **symplastic transport** in a plant root, represented by a purple arrow. **1**. AMF hypha **2**. Root **3**. Soil water **4**. Soil particles. **5**. Arbuscule **6**. Appressorium.

In soil, when water is transported by hyphae across an air gap, the soil solution likely contains nutrient ions that move to roots by diffusion and mass flow. When soil water content is very low, these nutrient supply paths are disrupted by the discontinuity of soil water films and plant nutrient deficiencies are common. AMF hyphal transport of water (outside of hyphal membranes) would not only supply water to roots during drought conditions but would also enable the movement of nutrient ions to roots in dry soils.

We believe our study is the first to explicitly test and observe water transport in hyphae by using isotopically labeled water to directly trace water flow from AMF to host plants. Our experimental method pairing H_2_^18^O and a fluorescent tracer provides strong evidence that AMF are able to bridge air gaps in soil and bring water to plants. In addition, our results indicate that water is transported to plants via an extracytoplasmic pathway in hyphae. Our findings have ramifications for the management of plant drought tolerance in the context of climate change. AMF symbioses are important actors in the maintenance of plant productivity when water is limited, making them essential not only in arid regions and semi-arid around the world, but also where short-term droughts occur^24^, especially as changing climatic conditions increase the occurrence of water-limiting conditions.

## Methods

### Experimental Set-up

*A. barbata* seeds gathered from the Hopland Research and Extension Center in Hopland, CA, were de-husked and sterilized in chlorine gas for 4 hours to kill any potential fungal pathogens on the surface and inside the seeds. Seeds were then germinated in Petri dishes on autoclaved filter paper and watered with autoclaved distilled water. The Petri dishes were placed in the dark at room temperature for two weeks. Seeds were de-husked and fumigated in chlorine gas for 4 hours.

Three two-week old *A. barbata* seedlings were planted in the ‘plant compartment’ of eighteen two-compartment microcosms (fig. 1). The plant compartment was separated from the ‘no-plant compartment’, by a 3.2 mm air gap. The purpose of the air gap was to prevent (or limit) liquid water from travelling passively between compartments. Both sides of the air gap had nylon mesh, either 18 μm, allowing hyphae but excluding roots, or 0.45 μm, excluding both hyphae and roots. A total of eighteen microcosms were used, twelve with 18 μm mesh and six with 0.45 μm mesh. The microcosms were made of laser cut 3.2 mm-thick acrylic panels glued together into single compartments; two compartments were screwed together tightly to make a microcosm (supplementary material figure 1). The plant compartment was 10×2.5×26.5 cm and packed with a sand-clay mixture, 1:1 by volume, to a 1.21 g/cm^3^ density (referred to herein as the ‘sand mix’). The no-plant compartment was 10×1×26.5 cm and packed with a soil-sand mixture, 1:1 by volume, to a 1.21 g/cm^3^ density (referred to as the ‘soil mix’). The sand and clay were washed three times in distilled water, air-dried, then autoclaved three times 48 hours apart. Soil (0 to 10 cm) was collected at the Hopland Research and Extension Center in Hopland, CA (38859.57840 N, 123804.04690 W) where *A. barbata* was the dominant vegetation type. The soil was then air-dried and sieved to 2 mm to remove large rocks and roots. The microcosms were covered with acrylic black panels on the outside to prevent growth of moss and algae. The no-plant compartment was covered at the top with AeraSeal breathable sealing films (152-68412, Spectrum Chemicals) to limit the dispersal of potential fungal pathogens from the soil mix in the no-plant compartment to the plant compartment.

In the plant compartment, the sand mix was inoculated with 26 g of whole inoculum of *R. intraradices* (accession number AZ243, International Culture Collection of (Vesicular) Arbuscular Mycorrhizal Fungi (INVAM), West Virginia University, Morgantown, WV). 20 g of the inoculum were mixed in the sand mix before packing, and the remaining 6 g were poured in a layer 3 cm from the top of the sand mix. In addition, 78 mg of autoclaved bone meal were mixed in the sand mix before packing, to encourage AMF growth and establishment. In the no-plant-compartment, 78 g of autoclaved bone meal were mixed in the soil mix before packing, to act as a bait for AMF to cross the air gap. The no-plant compartment was 1/4 of the volume of the plant compartment, therefore receiving four times as much bone meal as the plant compartment.

The microcosms were incubated in growth chambers in the Environmental Plant Isotope Chamber (EPIC) facility, located in the Oxford Tract Greenhouse at UC Berkeley, where environmental conditions can be monitored and controlled. Three chambers were used, with six microcosms in each, organized in a randomized fashion. Each microcosm was individually raised on two autoclaved metal bars to allow drainage and prevent water flow between microcosms. Chambers were thoroughly cleaned with a 10% bleach solution and a 70% ethanol solution before use. Each chamber had a fan with a cooling system to maintain temperature below 25°C. The fan also encouraged root growth due to the effect of simulated wind on shoots. Each compartment of the microcosms had three drain holes, 1 cm in diameter. Autoclaved glass wool was inserted into the drain holes to prevent soil from falling out, and 18 μm mesh was glued on the drain holes to prevent roots from growing out, while still allowing water to drain. Volumetric water content was monitored with electronic probes (EC-5, Decagon Services, Pullman, WA, USA) that measure the dielectric constant of the media. Two microcosms per chamber had probes, and the other four microcosms were assumed to have the same volumetric water content. Watering volumes were adjusted as needed to maintain a volumetric water content at approximately 17%. Both compartments of the microcosms were watered three times a week with autoclaved distilled water. 10 mL of filter-sterilized Rorison’s nutrient solution^25,26^ was added to the plant compartment (low P) and no-plant compartment (high P) once a week after distilled water. The plant compartment of microcosms with 0.45 μm mesh received twice as much nutrient solution as microcosms with 18 μm mesh, to make up for the nutrients plants could obtain via AMF in the no-plant compartment in microcosms with 18 μm mesh.

On day 1 of week 11 at 10 pm, 20 mL of water was injected using syringes with 15.2 cm long spinal tap needles into the no-plant-compartment (fig. 1). For the six microcosms with 18 μm mesh (the AMF-permitted ^18^O/dye microcosms, termed ‘+AMF’) and the six microcosms with 0.45 μm mesh (the AMF-excluded ^18^O/dye microcosms, termed ‘-AMF’), the water injected had a δ^18^O of +3000‰ and the fluorescent dye Lucifer Yellow carbohydrazide (LYCH, 0.01% w/v in water, MW 457). The ^18^O-labeled water and dye were added in order to trace the path of water from the no-plant compartment, through fungal hyphae crossing the air gap, to the plant roots. In the remaining six microcosms with 18 μm mesh (the AMF-permitted ^16^O/no dye microcosms, termed ‘^16^O’), the no-plant compartment was injected with water containing natural abundance water and no dye. The unenriched water we used was distilled water at natural abundance δ^18^O, that had been autoclaved three times for 30 minutes, 24 hours apart, to ensure it was free of fungal contaminants. The autoclaving process raised the δ^18^O value of the water from −12 to −8‰, but it remained in the natural abundance range. The ^16^O microcosms served to establish a baseline for the ^18^O/^16^O ratio and autofluorescence in plant and fungal tissues. The LYCH dye has been used in previous studies to investigate the path of hydraulically lifted water from plants to the soil through their mycorrhizal networks^27,28,29^, the reverse path of our experiment.

On day 2, 3, and 4 of week 11 from 6 am to 10 pm, a gallon-size plastic bag was placed over the shoots of each microcosm to collect transpired water. At 10 pm on each of the three days, the bags were carefully removed, sealed, and placed on ice overnight to allow the water to condense. The bags were weighed to measure the volume of water transpired, then the water was pipetted into tubes for isotopic analysis. We collected water each day for three days because we did not know how long it would take for water to be transported by AMF across the air gap and to be detectable in transpired water.

### Harvest and Sample Processing

On day 5 of week 11, all microcosms were destructively sampled. Shoots were cut at the base and into 1-inch pieces, dried at 60°C for 72 hours, and weighed for above ground biomass measurements. Roots were gently harvested and divided into several aliquots: (1) roots for staining with acid fuchsin were placed in distilled water, (2) roots for fluorescence microscopy were placed on wet kimwipes in Petri dishes and kept in the dark, (3) roots for molecular analysis were placed in cell release buffer^30^, and (4) roots for below ground biomass measurements were placed in paper envelopes and then dried.

The sand mix was collected and split up into several aliquots: (1) sand mix for volumetric water content was placed in 50 mL falcon tubes and stored at 4°C, then 10g was weighed, oven dried at 105°C until a stable weight was reached, and weighed again, (2) sand mix for molecular analysis was placed in a whirlpack bag, flash frozen in liquid nitrogen, and stored at −80°C, (3) sand mix for water extraction, was placed in a whirlpack bag, flash frozen in liquid nitrogen, and kept at −80°C, and (4) sand mix for nutrient measurements was placed in a whirlpack bag, flash frozen in liquid nitrogen, and stored at −80°C.

The soil mix was also collected and split up into several samples: (1) soil mix for gravimetric water content was placed in 50 mL falcon tubes and stored at 4°C, then 10g was weighed, oven dried at 105°C until a stable weight was reached, and weighed again, (2) soil mix for molecular analysis was placed in a whirlpack bag, flash frozen in liquid nitrogen, and stored at −80°C, (3) soil mix for water extraction was placed in a whirlpack bag, flash frozen in liquid nitrogen, and stored at −80°C, (4) soil mix for nutrient measurements was placed in a whirlpack bag, flash frozen in liquid nitrogen, and stored at −80°C, and (5) soil mix for hyphal extraction was placed in 50 mL falcon tubes and stored at 4°C; subsequently hyphae were removed using a dissecting microscope and tweezers.

Hyphae evident in the air gap were collected on the mesh facing the inside of the air gap using tweezers and scalpels and placed into the first tube of the DNeasy PowerSoil kit (Qiagen) for DNA extraction the same day.

## Microscopy

### Acid fuchsin

Roots were stained in acid fuchsin using a protocol modified from Habte & Osorio (2001)^31^. Roots were washed in distilled water, placed in a 10% KOH solution for 12 hours, then rinsed in distilled water. The roots were then placed in a 1% HCL solution for 12 hours, followed by a 0.01% acid fuchsin solution (85% 1M lactic acid, 5% glycerol in water) for 48 hours. Finally, the roots were put in a destaining solution (85% 1M lactic acid, 5% glycerol in water) for 48 hours. The stained roots were mounted on slides using the destaining solution and observed under both bright field and fluorescence (λ_ex_ 596 nm/ λ_em_ 615 nm).

### Fluorescent dye LYCH

For each microcosm, five 1-cm root segments were mounted on slides using a 50% glycerol solution in water. Fluorescence microscopy was conducted on the day of the harvest at the Biological Imaging Facility at UC Berkeley using the wavelengths λ_ex_ 428 nm/ λ_ex_ 536 nm that correspond to LYCH.

### Molecular Methods

For each microcosm, roots were washed twice in cell release buffer^30^ to remove microbial cells from their surfaces. Roots were then centrifuged to remove excess buffer, and ground in a tissue lyser with tungsten beads at 30 r/s for 20 min. DNA was extracted from 50 mg of ground roots for each microcosm using the DNeasy PowerPlant Pro Kit (Qiagen).

To extract AMF spores and hyphae from the soil mix, 4 g of homogenized soil mix were mixed with 50 mL of distilled water and 6 mL of hexametaphosphate solution (35g/L in water) and stirred for 30 min using a magnetic stirrer. The mixture was decanted through a 35 μm sieve and the spores and hyphae caught in the sieve were collected. This process was repeated twice for each microcosm to obtain enough spores and hyphae. DNA was extracted separately from spores and hyphae from the sand mix and from all of the air gap hyphae collected for each microcosm using the DNeasy PowerSoil kit (Qiagen).

DNA extracted from roots, soil mix spores and hyphae, and air gap hyphae was quantified by Quant-iT(tm) PicoGreen(tm) dsDNA Assay Kit (Invitrogen) and the concentrations were normalized. PCR was conducted on the normalized DNA samples using forward universal eukaryotic primer WANDA^32^ and reverse AMF-specific primer AML2^33^. This primer pair spans a variable 530-bp region in the SSU rRNA gene^34^. The PCR products were run on a gel to confirm the presence of DNA at 530 bp, then sequenced by Sanger sequencing at the UC Berkeley DNA Sequencing Facility. Sequencing results were compared to the Maarj*AM* database^35^ using the nucleotide BLAST function to confirm the presence of *R. intraradices* in roots, air gap, and soil mix. A sequence was considered a match for *R. intraradices* if query coverage and percent identity were both greater than 97%.

### Isotopic Analyses

Analyses of transpired water and water extracted from the soil mix and the sand mix were conducted at the Center for Stable Isotope Biogeochemistry (CSIB) at UC Berkeley.

Stable oxygen isotope composition of transpired water samples was determined by Isotope Ratio Infrared Spectroscopy (IRIS), using a L2140-i (Picarro Inc.) analyzer. Long-term external precision is ± 0.3‰.

Soil mix and sand mix water was extracted using a vacuum evaporation system and liquid nitrogen condensation trap. Stable oxygen isotope composition of soil mix and sand mix water extracts was measured by continuous flow (CF) using a Thermo Gas Bench II interfaced to a Thermo Delta V Plus mass spectrometer using a CO_2_-H_2_O equilibration method. In brief, 200 μL of water for both laboratory water standards and samples were pipetted into 10 mL glass vials (Exetainer®, Labco Ltd., UK) and quickly sealed. The vials were then purged with 0.2% CO_2_ in Helium and allowed to equilibrate at room temperature for 48 hours. The ^18^O/^16^O value of CO_2_ was then determined. Long-term external precision is ± 0.12‰.

### Statistical Analyses

Statistical analyses were conducted using R version 3.6.1^36^. A one-way analysis of variance (ANOVA) coupled with Fisher’s least-significant difference (LSD) test (package: agricolae) was used to differentiate means of ^18^O and volume of water transpired from different treatment groups.

### ^18^O Calculations

All the values used in the calculations below can be found in table 1 of the supplementary material.

#### Assumption #1

Water in the sand mix in the plant compartment of +AMF and -AMF microcosms that was directly accessible by roots initially had water with the same δ^18^O value as the water that was measured in the plant compartment of ^16^O microcosms at harvest, −1.53‰, since all microcosms received the same water throughout the experiment prior to the ^18^O-labeled water injection. This accounts for natural fractionation of water that may have occurred in the plant compartment (due to evaporation) after the plants were watered with natural abundance water with a δ^18^O value of −8‰.

#### Assumption #2

Water transported by AMF hyphae, or crossing the air gap as water vapor or droplets from the no-plant compartment, would have been diluted by water in the sand mix of the plant compartment before reaching roots. This dilution factor changed with time as the plant compartment water δ^18^O value increased from −1.53‰ at time = 0 to 162.19‰ at harvest. We assume that the dilution follows a linear model *y* = *ax* + *b*, where *x* represents time in days and *y* represents the δ^18^O value of the water in the plant compartment in ‰. We know from our measurements that *y*(0) = −1.53 and *y*(3.5) = 162.19, with the final harvest occurring at 3.5 days following the ^18^O-labeled water injection. Using the linear equation *y* = 46.78*x* −1.53, we calculated the following δ^18^O values: 45.25‰ for day 1, 92.03‰ for day 2, and 138.81‰ for day 3.

In equations 1-12 below, we define *x, y*, and *z* as follows:

***x*** = water directly taken up by roots and hyphae in the plant compartment

***y*** = water transported by AMF to the no-plant compartment taken up by roots

***z*** = water travelling via water vapor or droplet from no-plant to plant compartment taken up by roots

<u>Day 1</u>:

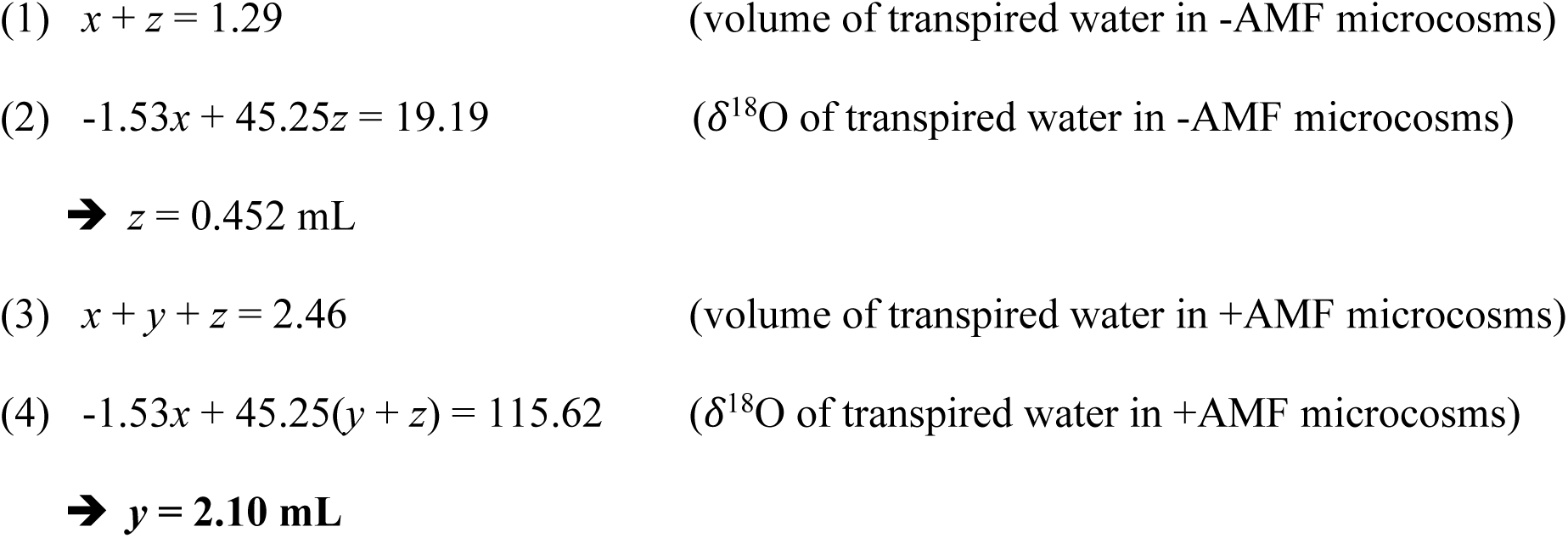

<u>Day 2</u>:

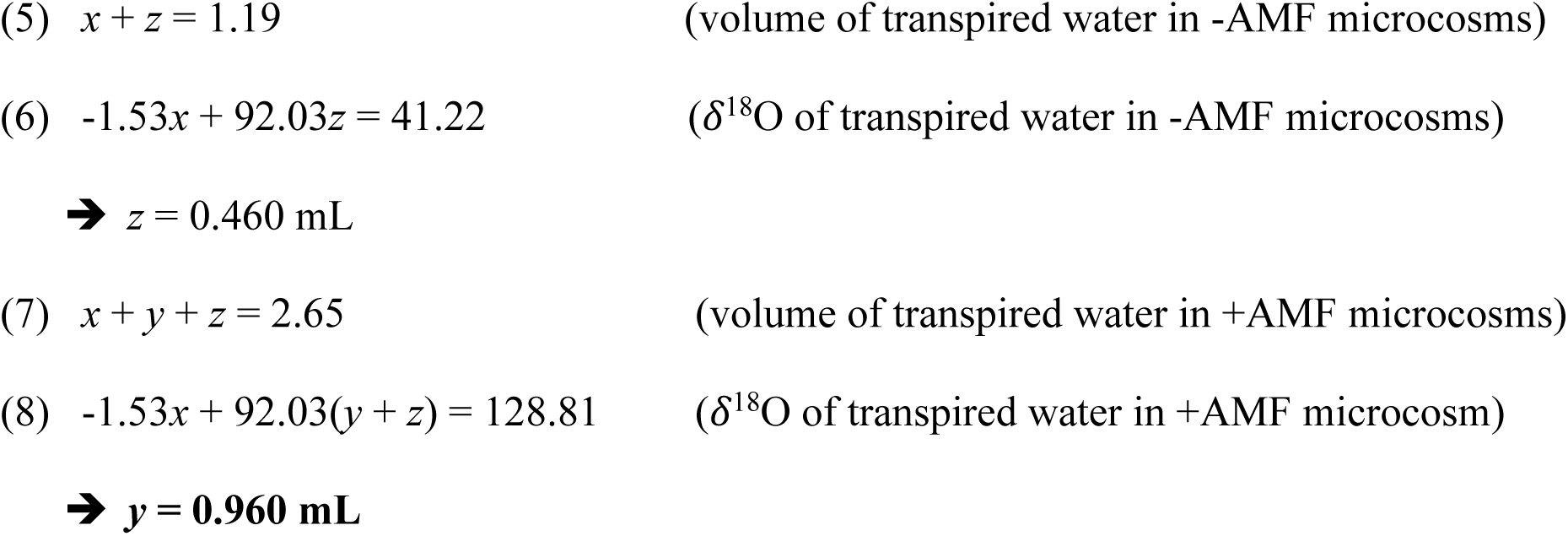

<u>Day 3</u>:

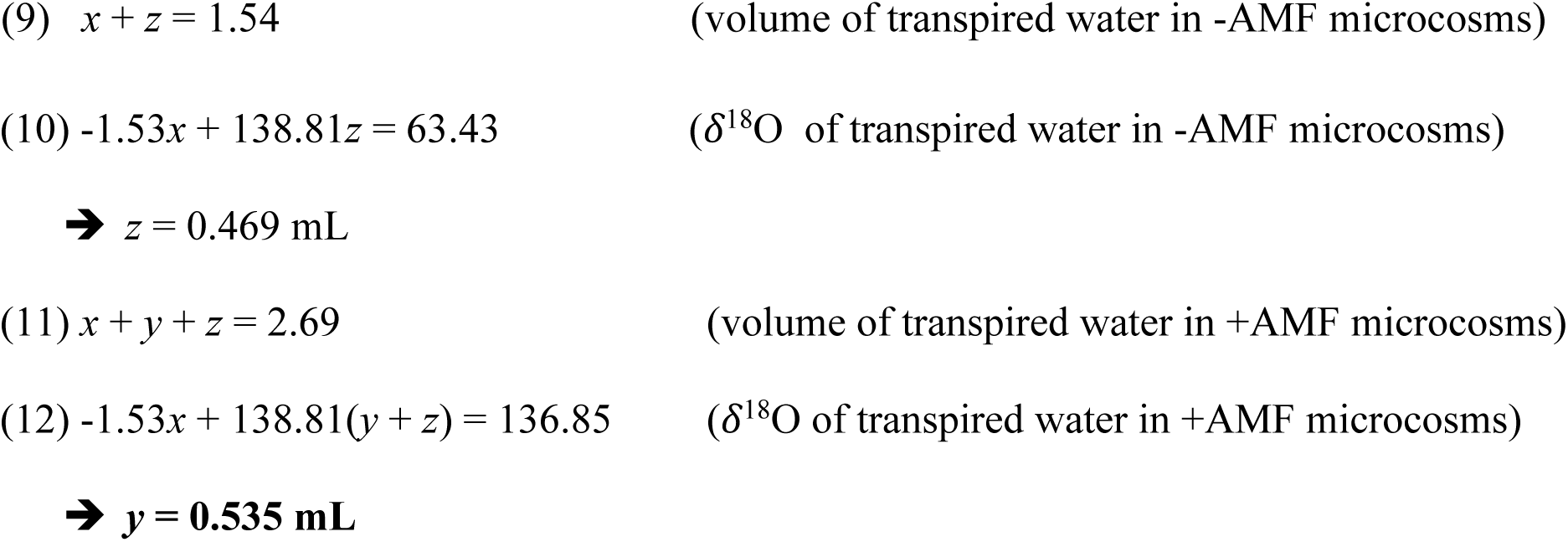

Over three days, +AMF plants transpired a total of 7.8 ±0.52 mL of water. This is 3.78 mL more than -AMF plants, which transpired a total of 4.02 ±0.79 mL. Based on the above calculations, the total AMF-transported water can account 46.2% of the total amount of water that was transpired by the +AMF plants over three days.

## Acknowledgements

The authors thank Dr. Thomas Bruns, Dr. John Taylor, Dr. Todd Dawson, Dr. Louise Glass, Dr. Angela Hodge and Dr. Greg Jedd for their thoughtful advice on the project, Dr. Denise Schichnes and Dr. Steven Ruzin for their guidance with microscopy, Christina Wistrom and Katerina Estera-Molina for their assistance with the greenhouse set up, and Hunter Jamison for his help with the isotope analyses.

## Authors Contribution

A.K. and M.K.F. designed the experiment with the assistance of J.A.H., D.J.H. and J.P-R. A.K., J.A.H and M.P.K performed the experiment with assistance from D.J.H. A.K., J.A.H, M.P.K, S.M. and M.K.F. analyzed the data with the assistance of L.J.F, D.J.H, and J.P-R. A.K. and M.K.F. drafted the manuscript. A.K., S.M., L.J.F., J.P-R., and M.K.F all contributed to the final manuscript.

## Data Availability

The data that supports the findings of this study are available from the corresponding author upon reasonable request.

