## Supplementary Material for "Routes to Roots: Direct Evidence of Water Transport by Arbuscular Mycorrhizal Fungi to Host Plants"

| | | "+AMF" | "-AMF" | "$^{16}$O" |
|---|---|---|---|---|
| transpired water | volume of transpired water at time = day 1 (mL) | 2.46 ±0.41 | 1.29 ±0.22 | 2.22 ±0.32 |
| | volume of transpired water at time = day 2 (mL) | 2.65 ±0.21 | 1.19 ±0.17 | 2.52 ±0.51 |
| | volume of transpired water at time = day 3 (mL) | 2.69 ±0.28 | 1.54 ±0.30 | 2.66 ±0.51 |
| | sum of volume of transpired water over three days (mL) | **7.80 ± 0.52** | **4.02 ±0.79** | 7.40 ±1.10 |
| | $\delta^{18}$O of transpired water at time = day 1 (‰) | 115.62 ±24.50 | 19.19 ±2.61 | 4.78 ±1.47 |
| | $\delta^{18}$O of transpired water at time = day 2 (‰) | 128.81 ±17.86 | 41.22 ±3.95 | 4.90 ±1.28 |
| | $\delta^{18}$O of transpired water at time = day 3 (‰) | 136.85 ±14.47 | 63.43 ±1.93 | 4.95 ±1.48 |
| | $\delta^{18}$O of transpired water over three days (‰) | **127.09 ±10.73** | **41.28 ± 5.75** | 4.88 ±0.78 |
| plant compartment | mass of dry sand-clay mixture (g) | 745.1 | 745.1 | 745.1 |
| | gravimetric water content of sand-clay mixture at harvest (%) | 16.5 ±0.6 | 15.9 ±1.0 | 16.1 ±0.5 |
| | volume of water in sand-clay mixture at harvest (mL) | 122.83 ±4.24 | 116.12 ±3.42 | 117.07 ±7.70 |
| | $\delta^{18}$O of water in sand-clay mixture at harvest (‰) | **162.19 ±10.80** | **80.70 ±12.51** | -1.53 ±1.97 |
| | Above ground biomass at harvest (mg) | 559.2 ±64.5 | 535 ±49.7 | 769.2 ±76.6 |
| | Below ground biomass at harvest (mg) | 805.3 ±80.5 | 801.9 ±75.2 | 814.5 ±88.8 |
| | Above ground biomass %C | 37.73 ±0.42 | 29.73 ±7.21 | 37.03 ±0.60 |
| | Above ground biomass %N | 0.87 ±0.02 | 0.60 ±0.16 | 0.81 ±0.09 |
| | Above ground biomass C:N | 42.62 ±1.37 | 50.11 ±2.20 | 49.00 ±6.79 |
| | Above ground biomass %P | **0.29 ±0.05** | **0.12 ±0.012** | 0.25 ±0.012 |
| no-plant compartment | mass of dry soil-sand mixture (g) | 230.0 | 230.0 | 230.0 |
| | gravimetric water content of soil-sand mixture at harvest (%) | 11.6 ±0.3 | 11.1 ±1.1 | 9.9 ±0.4 |
| | volume of water in soil-sand mixture at harvest (mL) | 25.63 ±2.45 | 24.79 ±0.62 | 25.77 ±0.84 |
| | $\delta^{18}$O of water added at t=0 (‰) | 3000.00 | 3000.00 | -8.00 |
| | $\delta^{18}$O of water in soil-sand mixture at harvest (‰) | 300.75 ±50.88 | N/A | N/A |

**Table 1**: Data used in statistical analyses and $^{18}$O calculations. The values that appear in the main text are in **bold**. Means ± standard error are averages of six microcosms, each having three plants (one-way ANOVA & Fisher LSD test).

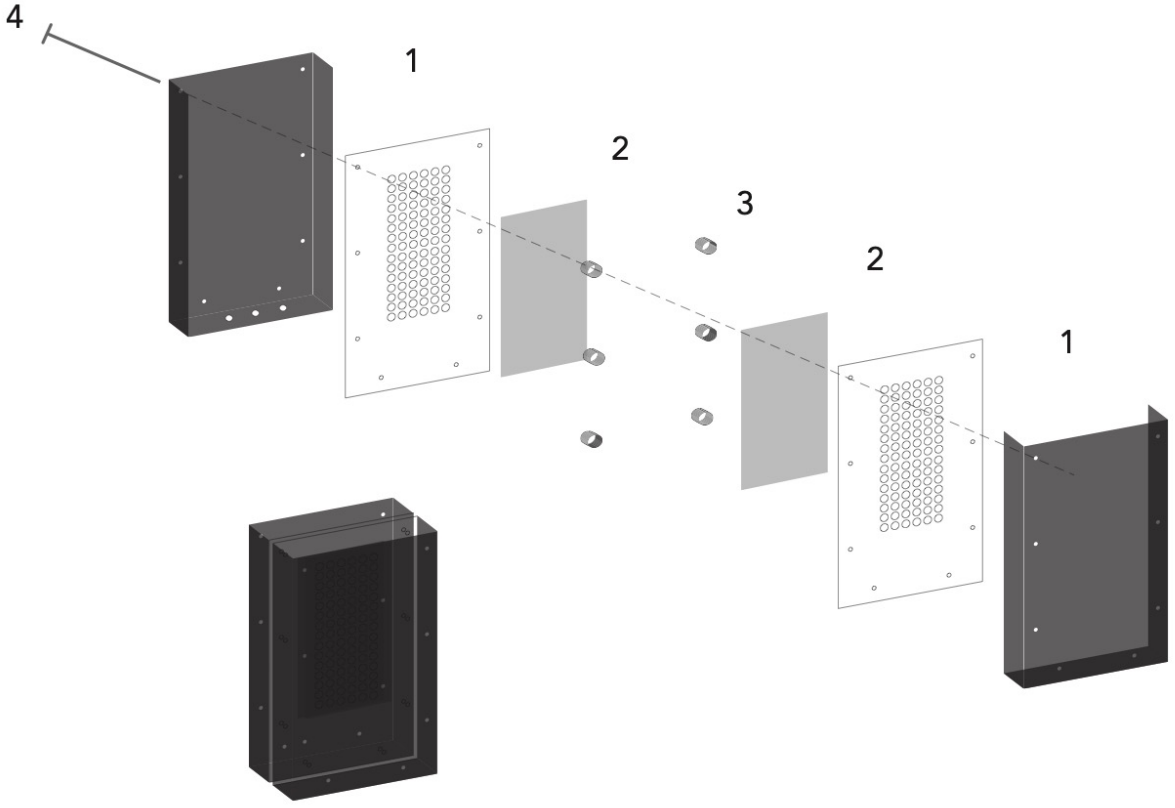


**Figure 1**: Assembly of a microcosm. **1.** laser cut acrylic panels, **2.** nylon mesh, **3.** acrylic washers, and **4.** metal screws.

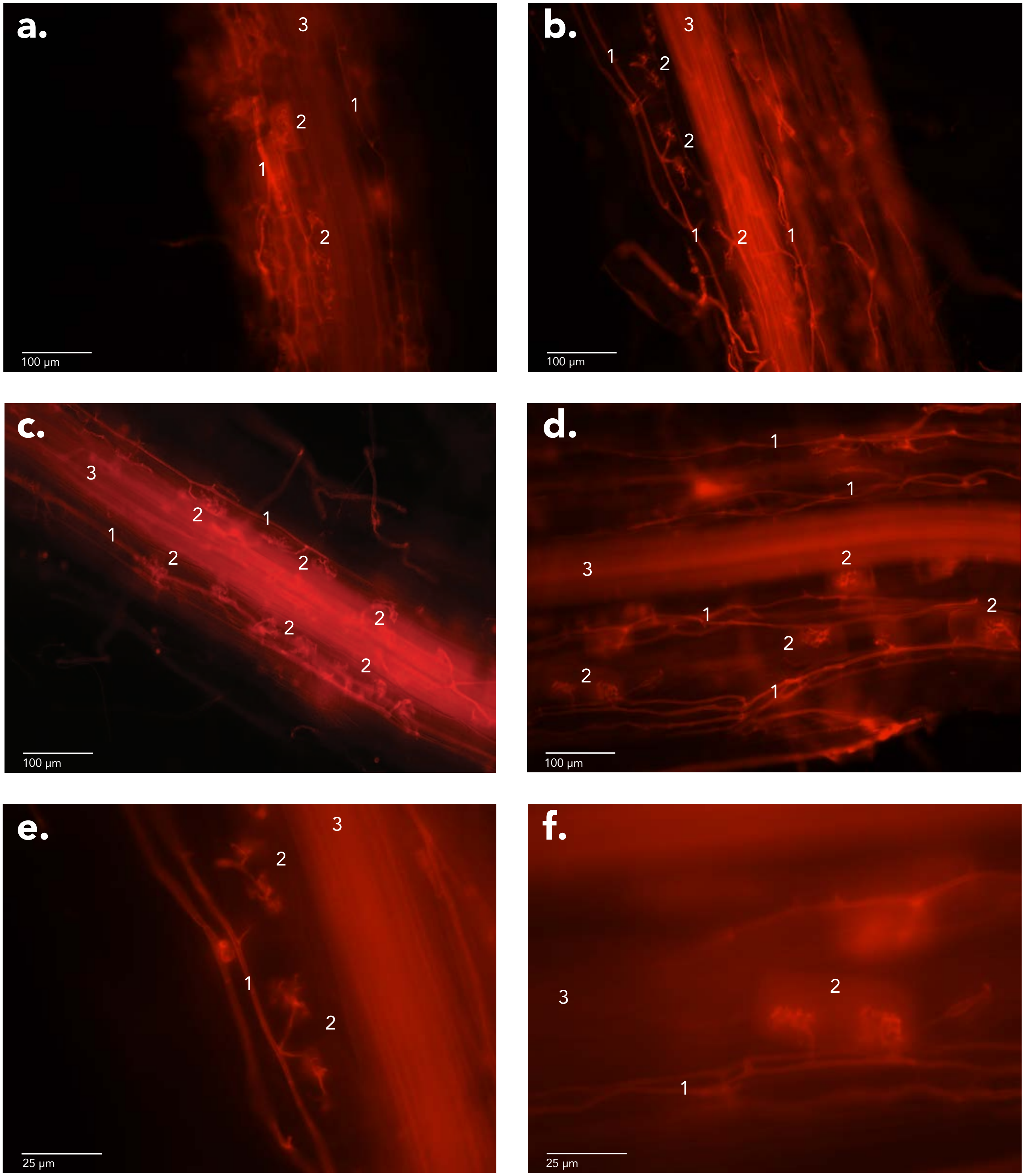


**Figure 2**: *Avena barbata* roots dyed with acid fuchsin showing AMF structures. **a-f.** Fluorescence images at AMF wavelengths (**$\lambda_{ex}$** 596 nm; **$\lambda_{em}$** 615 nm).**1.** Hyphae. **2.** Arbuscules. **3.** Root.
